# Sleep affects low-gamma range effective cortical connectivity for 40-Hz auditory steady-state responses

**DOI:** 10.64898/2026.02.13.705834

**Authors:** Anna Leśniewska, Urszula Górska-Klimowska, Miroslaw Wyczesany, Małgorzata Hołda, Bartłomiej Chojnacki, Marek Binder

## Abstract

The 40-Hz auditory steady-state response (40-Hz ASSR) is a sensitive marker of changes in arousal level, which has been reported to decrease during slow-wave sleep. However, sleep-related changes in directional connectivity during 40-Hz ASSR across cortical networks remain underexplored. In this study, we examined how wakefulness, NREM (N1, N2, N3) and REM sleep affect the direction and extent of neural signal propagation. EEG data during periodic 40-Hz auditory stimulation were collected during an overnight study from 29 normal-hearing human subjects (including 16 females). A source analysis was implemented to locate cortical activity, and effective connectivity was assessed with the Directed Transfer Function (DTF) in the low-gamma band (37-43 Hz). We focused on the connections between auditory cortical regions, prefrontal and temporo-parietal associative cortices. We hypothesized that: 1) feedback connections from associative to primary auditory areas will be the most affected by the arousal state changes; 2) associative reciprocal connectivity between prefrontal and temporo-parietal regions will display gradual connectivity reduction with increasing NREM sleep depth, with partial restoration during REM sleep.

Our results showed that feedforward rather than feedback connectivity was most strongly disrupted during sleep, particularly in NREM N2 and N3 stages, contradicting our first hypothesis. The second hypothesis was supported: reciprocal connectivity between prefrontal and parietal associative cortices significantly decreased with sleep depth. Overall, our findings suggest that reduced cortical propagation of 40-Hz ASSR related neuronal signals during sleep primarily reflects a breakdown in bottom-up signal transmission, and a parallel weakening of reciprocal prefrontal-parietal coupling.

## Introduction

Sleep is a dynamic state during which the brain’s responsiveness to external stimuli, including auditory inputs, is markedly modulated. Different sleep stages—N1, N2, N3 (NREM), and REM—exert distinct effects on auditory processing (Strauss et al., 2015; Tononi & Massimini, 2008). In addition to thalamic gating phenomenon, sleep appears to disrupt long-range cortico-cortical communication and limiting auditory information integration across distributed networks (Boly et al., 2012; Massimini et al., 2005).

The 40-Hz auditory steady-state response (ASSR), an oscillatory EEG response evoked by periodic sounds, is highly sensitive to global arousal levels fluctuations. It shows significant suppression during NREM sleep stages N2 and N3 (Górska & Binder, 2019; Suzuki et al., 1994), with only partial recovery during REM sleep (Cohen et al., 1991) and returning to wakefulness levels upon awakening (Picton et al., 2003). This attenuation involves reductions in both amplitude and intertrial phase coherence (Górska & Binder, 2019) and correlates with decreased activity in auditory associative areas (Issa & Wang, 2011). On the other hand, the patterns of cortical propagation of 40-Hz ASSR across sleep stages remain less understood.

Our study investigates this issue using effective connectivity measures, focusing on the regions that are critical for auditory processing. They include the primary auditory cortex (PAC) and the superior temporal gyrus (STG), both implicated in 40-Hz ASSR generation (Herdman et al., 2002; Spencer, 2012). During wakefulness, PAC shows strong, right-lateralized activation (Ross et al., 2005), but during sleep, its activity diminishes due to reduced thalamic input, especially during NREM sleep (Edeline et al., 2000) and phasic REM (Wehrle et al., 2007). The STG, involved in hierarchical auditory processing, supports multisensory integration (Kuśmierek & Rauschecker, 2009; Medalla & Barbas, 2014; Poremba et al., 2004) and interacts reciprocally with primary auditory and prefrontal regions, such as the anterior prefrontal cortex (aPFC) and dorsolateral prefrontal cortex (dlPFC) (Plakke & Romanski, 2014). These connections may support the integration of complex auditory information and cognitive monitoring, also during sleep (Pugh et al., 1996; Tzourio et al., 1997). Additionally, regions like the temporoparietal junction (TPJ) and posterior inferior parietal lobule (pIPL) are involved in bottom-up attention (Astafiev et al., 2006; Corbetta & Shulman, 2002) and multisensory integration (Zvyagintsev et al., 2013), and creating the sensation of perceptual awareness Gillebert et al., 2011; Mesulam, 1998). To investigate directed connectivity across sleep stages, we focused on three connection types: (i) feedforward and (ii) feedback connections between lower- and higher-order auditory regions, and (iii) reciprocal connections between associative cortices.

We have hypothesized that (1) the reduction in 40-Hz ASSR propagation during sleep is associated with changes in both auditory feedforward and feedback connectivity, with feedback connectivity being especially vulnerable during NREM N2 and N3 stages. This is supported by prior studies indicating that loss of conscious auditory perception is associated with impaired hierarchical processing including cortical regions in auditory cortex (Boly et al., 2011; Sanders et al., 2018). While many reports emphasize disrupted feedback connectivity in unconscious states (Boly et al., 2011; Lamme & Roelfsema, 2000), impairments in feedforward pathways with proceeding loss of consciousness have also been observed (Allen et al., 2020; Railo et al., 2011; Sanders et al., 2018). We have further hypothesized that (2) reciprocal connectivity between associative regions in prefrontal and parietal cortices will progressively weaken from wakefulness to NREM N2 and N3 sleep and partially recover during REM sleep. This is motivated by findings that associative areas are key to conscious processing (Naghavi & Nyberg, 2005; Sarter et al., 2001), and that functional connectivity between them decreases with increasing sleep depth (Fernandez Guerrero & Achermann, 2018; Sämann et al., 2010). Importantly, recent work suggests these connections are mutually reinforcing rather than strictly hierarchical (Nee, 2021), underscoring the relevance of their bi-directional dynamics across consciousness states.

## Materials and Methods

### Participants

The experiment was conducted in accordance with the directive of the Declaration of Helsinki (1975, revised in 2000) and approved by the Research Ethics Committee at the Institute of Psychology, Jagiellonian University. Before beginning the study, all participants provided written informed consent and confirmed that they were currently drug-free, had no history of substance dependence, and had no documented neurological or psychiatric disorders. The sample consisted of 29 participants (16 females, 55.2%) aged 19 to 55 years (mean = 29.8, SD = 9.9 years). Prior to the study, participants were instructed to restrict their sleep to 3-5 hours the night before to increase sleep pressure and facilitate falling asleep in an unfamiliar environment. All participants underwent hearing screening using the Titan device v. 3.4.1 (Interacoustics A/S, Middelfart, DK), testing integrity of the inner ear with otoacoustic emissions and integrity of the auditory pathway with auditory brainstem responses. Only participants who passed the screening process (i.e. those who met at least one criterion with a ‘pass’ result) were allowed to participate in the main experiment.

### Auditory stimulation protocol

The stimuli consisted of 500-ms long click trains presented at 40 Hz, with inter-stimuli intervals randomly set between 700 and 1000 ms (step 100 ms). Each participant received 100 repetitions of the stimulus, presented binaurally at 60 dB. Acoustic calibration was performed by playing the stimulus through the complete measurement system and recording the SPL using in-ear microphones embedded in a B&K Head and Torso Simulator HATS 4128-C (Hottinger Brüel & Kjaer GmbH, Darmstadt, DE). The details of the sound pressure level calibration procedure are described in the Supplementary Material (Supplemental Text S1)

Three other types of auditory stimulus, which were not analyzed in this study, were presented within the same session, which thus lasted about 10 minutes. Each participant underwent 3–8 sessions during their overnight stay in the lab. Auditory stimuli were prepared in MATLAB environment (The MathWorks, Inc., Natick, MA, USA) and delivered using ER-3C insert earphones (Etymotic Research, Elk Grove Village, IL, USA) and a headphone amplifier Millenium HP1.

### Apparatus

EEG recordings were carried out using 64-channel Active Two system (BioSemi Active Two). Electrodes were positioned on the scalp with two additional reference electrodes placed on the mastoid. To ensure compatibility with the 10-20 system, Standard Electro-Cap electrodes (Electro-Cap International Inc., Eaton, USA). A baseline EEG signal was recorded for two minutes after the electrodes were mounted. The data were sampled at a rate of 1024 Hz. The audio signal was recorded concurrently with EEG data using Analog Input Box (Biosemi, Amsterdam, NL) and stored in one dataset. Additionally, polysomnography was conducted using the Grass Technologies Comet PSG system (Natus Medical Inc., Pleasanton, CA, USA), with gold-plated cup electrodes applied on the face and scalp. This setup included three EEG channels (F4, C4, and O2, referenced to Cz), two EOG channels positioned in the outer canthi of the left and right eye (referenced to M2), and three EMG channels (recorded as a bipolar derivation) to monitor the chin muscle tone. The EEG channels were repositioned 1 cm back and to the right from their original positions to avoid overlapping with the corresponding channels on the EEG cap.

### Procedure

The experiment took place overnight in the Sleep Psychology Laboratory, Institute of Psychology, Jagiellonian University. Participants slept in a separate, air-conditioned room with the lights switched off. Each participant slept for no more than 7.5 hours during the experimental night. The procedure began about 9 pm with the auditory screening, and after obtaining the passing result, participants were given time for a bedtime routine, and EEG and PSG electrodes were then placed on their scalp and face. Stimulus presentation was controlled by Presentation® software (Version 23.0, Neurobehavioral Systems, Inc., Berkeley, CA). The actual experiment with auditory stimulation and EEG/PSG data acquisition was conducted during sleep and during wakefulness before falling asleep (and after waking up, but the latter was not analyzed in this work). During wakefulness sessions, participants were asked to sit comfortably with their eyes open (EO) and eyes closed (EC) (the latter, again, was not analyzed here), allowing their thoughts to flow spontaneously without focusing on the auditory stimulation or their thoughts. During sleep sessions, auditory stimuli were presented 3 to 8 times to capture recordings across all sleep stages.

### Data analysis

#### Sleep staging

The Sleep staging solution tool in Brain Vision Analyzer 2.2 software (Brain Products, Gilching, DE) was used to assess sleep stages by the two authors (AL, UGK). The classification of sleep stages in the EEG data was cross-verified with hypnograms based on PSG recordings visually scored in 30-s epochs by another author (MH), thus ensuring that stages were correctly identified by multiple independent examiners. The sleep stages considered for subsequent analysis were N1, N2, N3 NREM, and REM. Sleep staging adhered to the guidelines of the American Academy of Sleep Medicine (AASM) (Berry et al., 2017).

#### Preprocessing of EEG data

Preprocessing was carried out using the Atlantis connectivity toolbox (Wyczesany et al., 2025). The EEG signal was then filtered using windowed sinc linear phase FIR filters, with a 1 Hz high-pass filter (order: 3300) and a 48 Hz low-pass filter (order: 1690). Next, the data was segmented into epochs with one second duration, beginning at the stimulus onset with segments demeaning. Next, channels were screened using the interquartile range (IQR) method and marked as bad if their variance fell outside the Q1/Q3 ± 4 x IQR range.

Surviving original channels were re-referenced to the P9 and P10 channels. Trial-based artifact rejection consisted of extreme outliers removal based on variance (IQR threshold of ± 6 x IQR), maximum trial voltage difference (< 400 μV), and identifying muscle artifacts. The latter were characterized by elevated spectral power (*z* = 20) in the 35-47 Hz range, where z-scores represent how many standard deviations a component’s spectral power deviates from the mean across all components. The remaining signal was decomposed using the fast-ICA method with deflation enabled. IC time courses were further screened for artifacts and removed if they exhibited trial excess variance (*z* > 20 for a single IC or *z* > 6 on at least 6 ICs).

The pruned signal was decomposed again using ICA, repeated 22 times. For each decomposition, components were classified using a pre-trained model installed in the Atlantis tool that considered features such as topography, spectral power, pre/post stimulus variance, and signal kurtosis to distinguish between brain and non-brain ICs. The best ICA decomposition was chosen based on the number of brain components and artifact contamination metrics. Signal components from the selected ICA realization were localized using their topographies with the minimum norm estimation (MNE) method (Hämäläinen & Ilmoniemi, 1994) based on a standard, single-shell Montreal Neurological Institute (MNI) template head model (Colin27) with depth weighting parameter d set to 0.5 (further details of the analytic procedure can be found in Adamczyk and Wyczesany, 2023).

Region of interest (ROI) signals were reconstructed as the sum of IC signals in the respective (closest) source dipole. Sources were obtained as the product of a particular IC time course and respective weight components separately for all spatial directions. The scalar value of the ROI signal was represented by the first principal component derived from the three spatial vectors. To control spurious correlations between estimated sources, leakage correction for multivariate data was applied (Colclough et al., 2015). The ROIs and their coordinates were derived from the MNI atlas and are listed in Table 1.

**Table 1.** Selected regions of interest with the Montreal Neurological Institute (MNI) coordinates and responding Brodmann’s Areas. The regions of interest were located bilaterally (L = Left, R = Right) in prefrontal cortex (anterior prefrontal cortex, aPFC and dorsolateral prefrontal cortex, dlPFC), auditory cortex (primary auditory cortex, AC and secondary auditory cortex, STG) and posterior associative cortex (temporoparietal junction, TPJ and angular gyrus/ posterior intraparietal lobule pIPL).

| Selected regions of interest (ROIs) |  |  |  |  |
| --- | --- | --- | --- | --- |
|  | Abbreviation | Region | MNI coordinates | Brodmann's Area |
| Left Primary Auditory Cortex | LPAC | Auditory cortex | -50 -21 7 | 41 |
| Right Primary Auditory Cortex | RPAC | Auditory cortex | 50 -21 17 | 41 |
| Left Superior Temporal Gyrus | LSTG | Auditory cortex | -64 -35 9 | 22 |
| Right Superior Temporal Gyrus | RSTG | Auditory cortex | 64 -35 12 | 22 |
| Left Anterior PFC | LaPFC | Prefrontal cortex | -35 45 30 | 10 |
| Right Anterior PFC | RaPFC | Prefrontal cortex | 32 45 30 | 10 |
| Left Dorsolateral PFC | LdIPFC | Prefrontal cortex | -43 38 12 | 46 |
| Right Dorsolateral PFC | RdIPFC | Prefrontal cortex | 43 38 12 | 46 |
| Left Temporoparietal Junction | LTPJ | Parietal cortex | -45 -50 22 | 39 |
| Right Temporoparietal Junction | RTPJ | Parietal cortex | 45 -50 22 | 39 |
| Left Intraparietal Lobule (posterior) | LpIPL | Parietal cortex | -39 -75 33 | 39/19 |
| Right Intraparietal Lobule (posterior) | RpIPL | Parietal cortex | 39 -75 33 | 39/19 |

#### Effective connectivity analysis

Connectivity between the chosen ROIs was estimated for low gamma (37-40 Hz) frequency band using the directed transfer function (DTF; Kaminski & Blinowska, 2017), a multivariate method based on Granger causality assumptions. Connectivity values were estimated for the 0-500 ms epochs corresponding to the ongoing stimulus time windows.

DTF offers several advantages relevant to the aims of our study that include direct assessment of connectivity in the low-gamma band (∼40 Hz): including directionality inference, simultaneous modeling of multiple cortical interactions, and frequency-specific analysis.

In accordance with the study’s aims, directed connections between selected ROIs were segregated into three functional connectivity groups: auditory feedforward, auditory feedback, and associative connections. Feedforward connections were defined as ipsilateral projections from the primary auditory cortex (PAC) and superior temporal gyrus (STG) to higher-order areas, including anterior prefrontal cortex (aPFC), dorsolateral prefrontal cortex (dlPFC), temporoparietal junction (TPJ), and posterior inferior parietal lobule (pIPL), while feedback connections were operationalized as projections in the reverse direction. In order to maintain a clear assessment of hierarchical organization, the analysis was restricted to ipsilateral pathways. Ipsilateral connections predominantly support hierarchical, sequential processing (Markov et al., 2014) whereas contralateral pathways are more often involved in interhemispheric integration rather than strict feedforward or feedback transmission. Accordingly, in assessing reciprocal connectivity between associative hubs (dlPFC, TPJ, and pIPL), both ipsilateral and contralateral directional pathways were analyzed. This approach allowed for the characterization not only of localized hierarchical dynamics but also of long-range interhemispheric processes that contribute to perceptual and cognitive integration (Sousa et al., 2019).

#### Statistical testing

Statistical analyses were conducted using R Statistical Software (v.4.2.2; R Core Team, 2023). To investigate the effects of arousal changes on directed connectivity values derived from the Directed Transfer Function (DTF) under 40-Hz ASSR entrainment, we employed linear mixed-effects models (LMEMs) for modeling individual connections, using the lme4 package (v.1.1-33; Bates et al., 2015). Estimated marginal means (EMMs) were computed to perform pairwise comparisons of connectivity estimates between wakefulness and sleep stages for each directed connection.

To account for repeated measures across participants, each model included arousal stage as a fixed effect and subject (id) as a random intercept. The optimizer “bobyqa” was applied to improve model convergence. However, for 7 of the 48 connections, model fitting indicated singularity (as detected by the isSingular() function), suggesting issues with the variance component for the random intercept. For these connections, simplified linear models including only fixed effects were used instead. More information about steps included in the statistical modelling can be found in the supplementary.

Arousal stages were coded as a factor variable with wakefulness (W) as the reference level. Pairwise comparisons between W and each sleep stage contrasts were conducted using estimated marginal means with emmeans package (v.1.8.6; Lenth, 2023), with effect sizes (Cohen’s d) computed using the lme.dscore function from the EMAtools package (v.0.1.4; Kleiman, 2021). For each direction in the contrast, we provide a normalized estimated marginal mean difference (B), which represents the effect of sleep stage on DTF connectivity value and facilitates direct comparisons of effect sizes between different arousal stages contrasts. Confidence intervals (95% CI) for these estimates indicate the range within which the true effect is likely to fall. We applied FDR correction across all individual connection results in each contrast.

## Results

The results presented below illustrate brain patterns of disrupted effective connectivity related to 40-Hz ASSR in contrasts between sleep stages and wakefulness. The connections shown are those whose *p*-values survived the False Discovery Rate (FDR) correction (*p_FDR_* < 0.05) and differentiate the contrasted arousal states in low-gamma band (Figure 1 and Tables 2.A-2.D).

**Figure 1.**
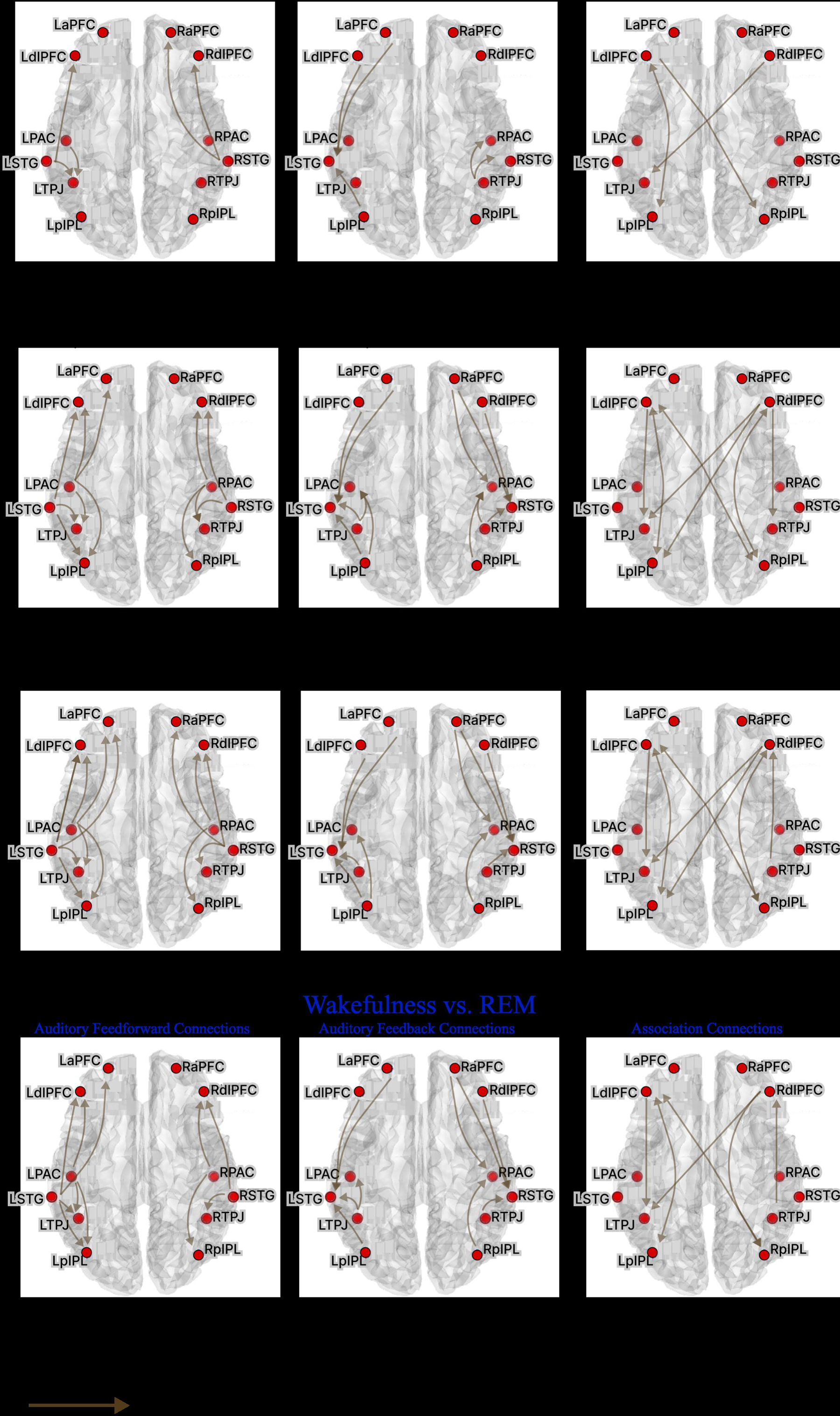
*Contrasts of DTF between wakefulness with sleep stages within auditory feedback, auditory feedforward and associative connection networks.* Effective connectivity maps representing results of contrasting the DTF scores between wakefulness with sleep stages (NREM N1, N2, N3, and REM) within three functional networks: auditory feedforward connections (left column), auditory feedback connections (middle column), and associative connections (right column). Each panel shows only significant reductions, indicated by arrows, in cortico-cortical effective connectivity during sleep (p<0.05, FDR-corrected) in the low-gamma band (37–43 Hz). The arrows represent directed connections with significantly reduced effective connectivity in the sleep stage as contrasted with wakefulness. (LaPFC – Left anterior Prefrontal Cortex, RaPFC – Right anterior Prefrontal Cortex, LdlPFC – Left dorsolateral Prefrontal Cortex, RdlPFC – Right dorsolateral Prefrontal Cortex, LPAC – Left Primary Auditory Cortex, RPAC – Right Primary Auditory Cortex, LSTG – Left Supratemporal Gyrus, RSTG – Right Supratemporal Gyrus, LTPJ – Left Temporoparietal Junction, RTPJ – Right Temporoparietal Junction, LpIPL – Left posterior Intraparietal Lobule, RpIPL – Right posterior Intraparietal Lobule).

**Table 2.**
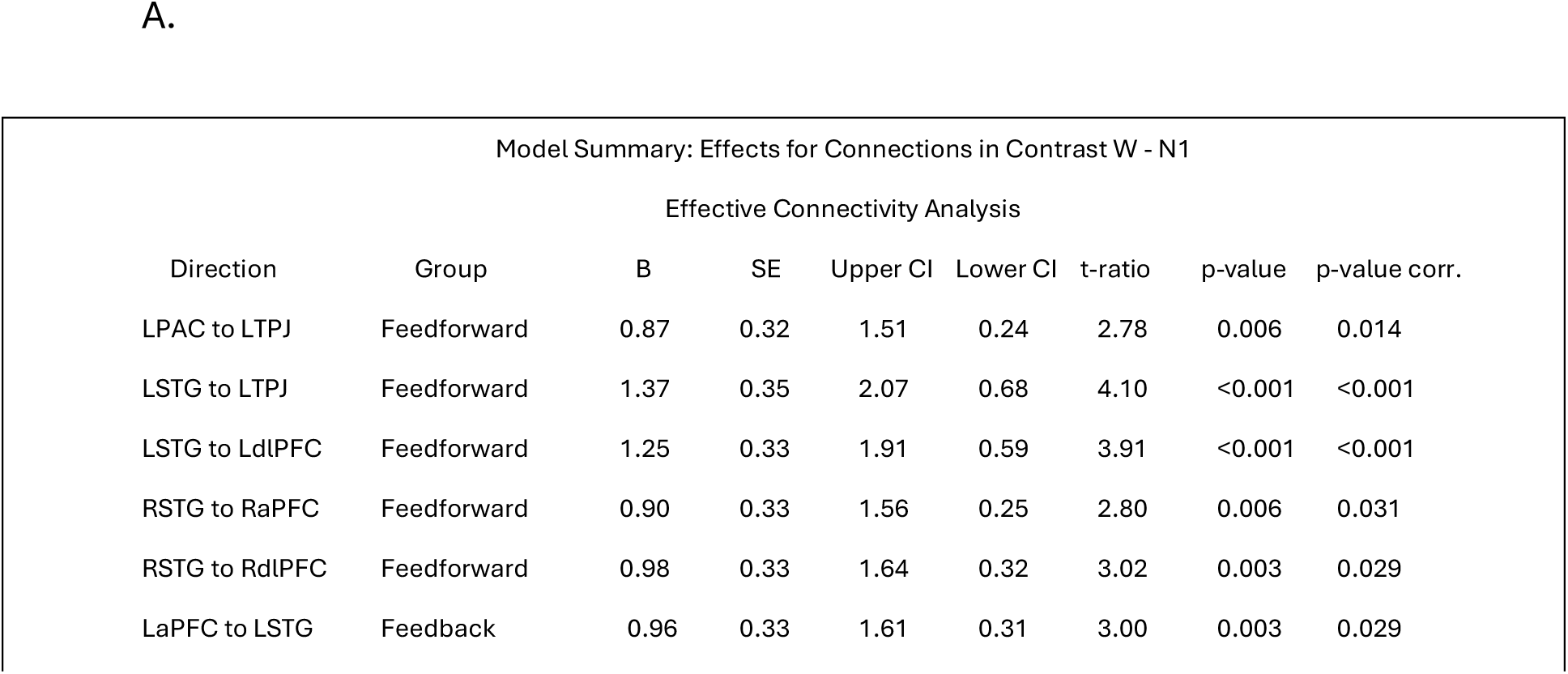
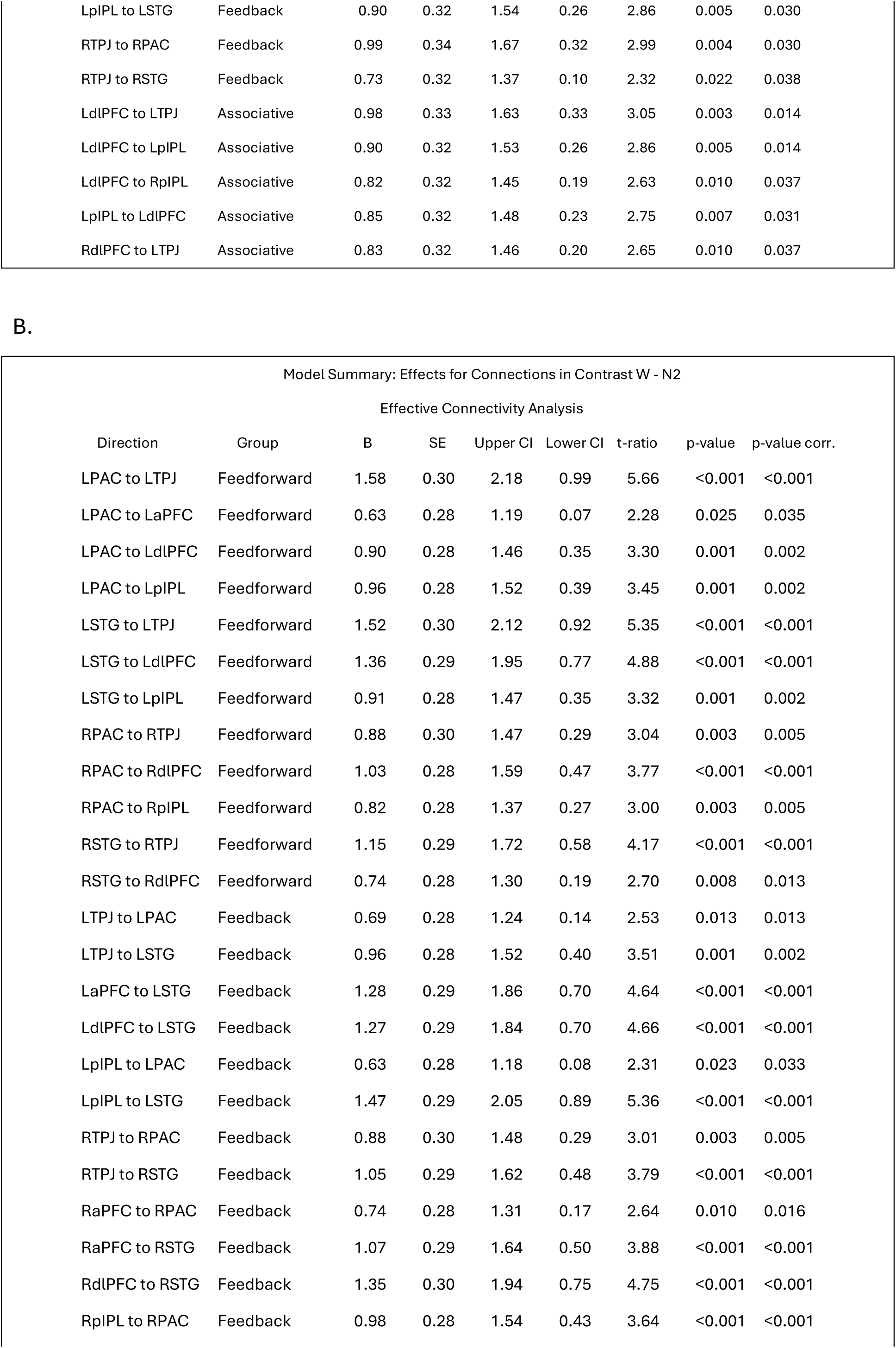
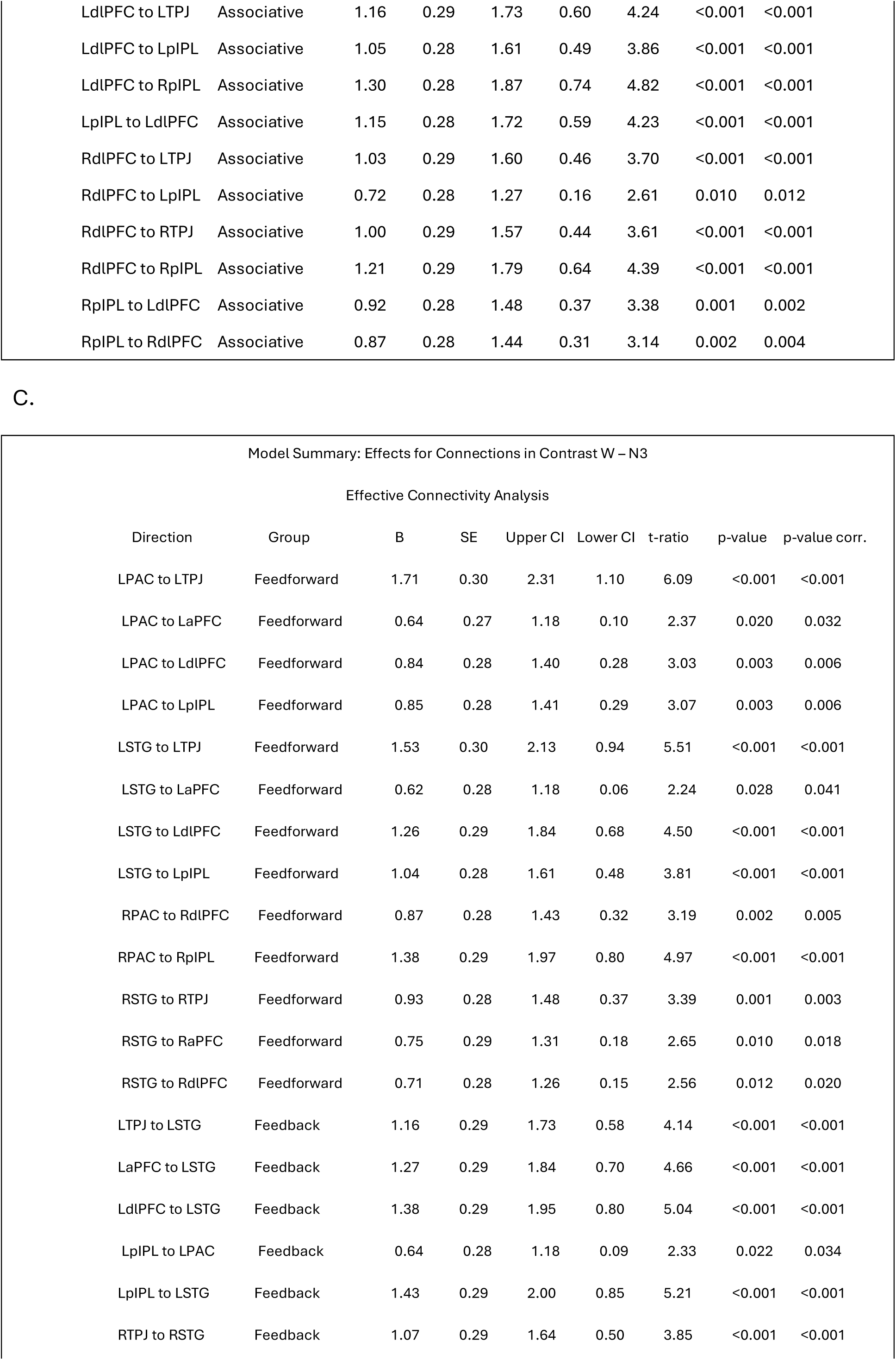
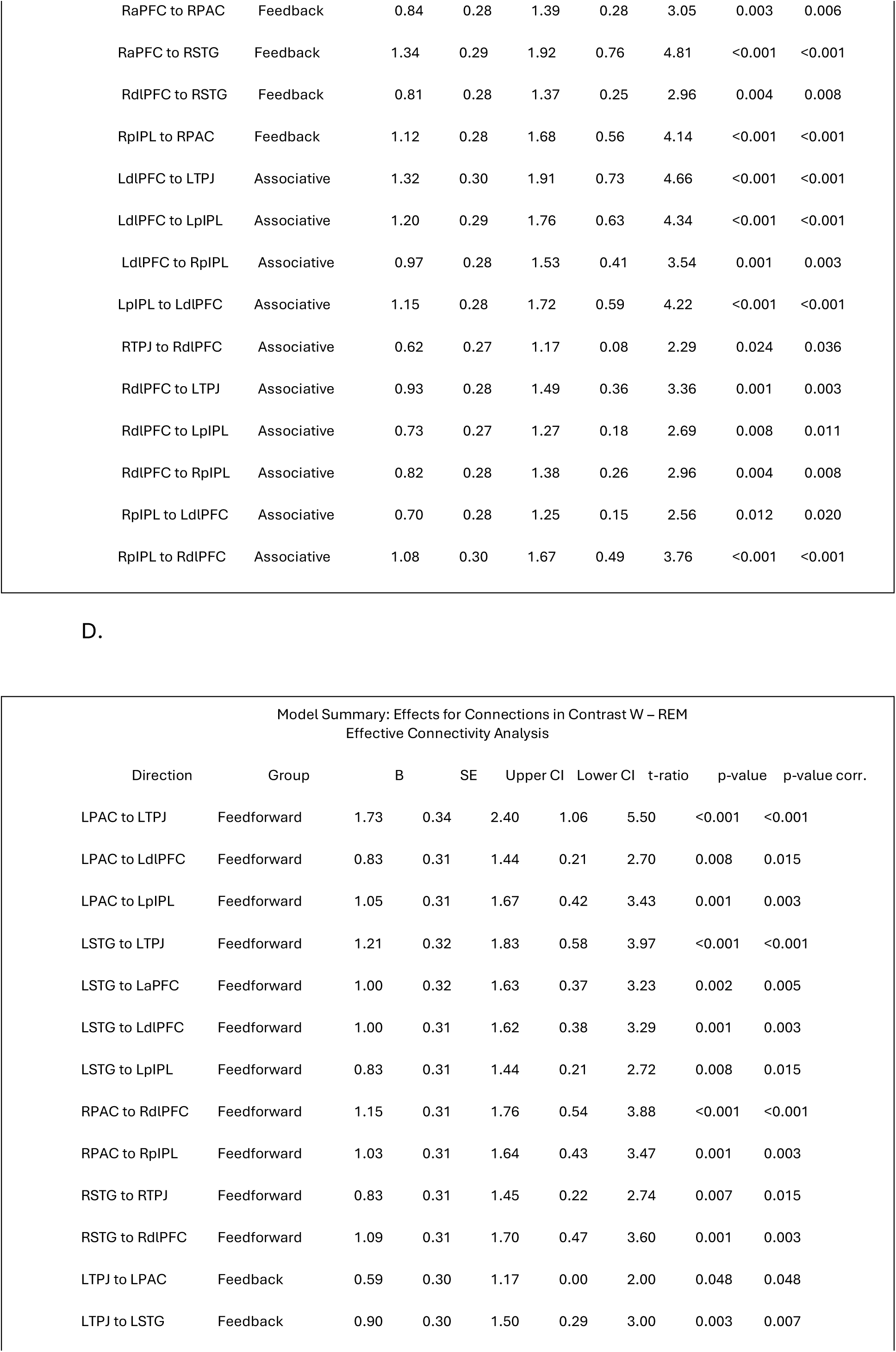
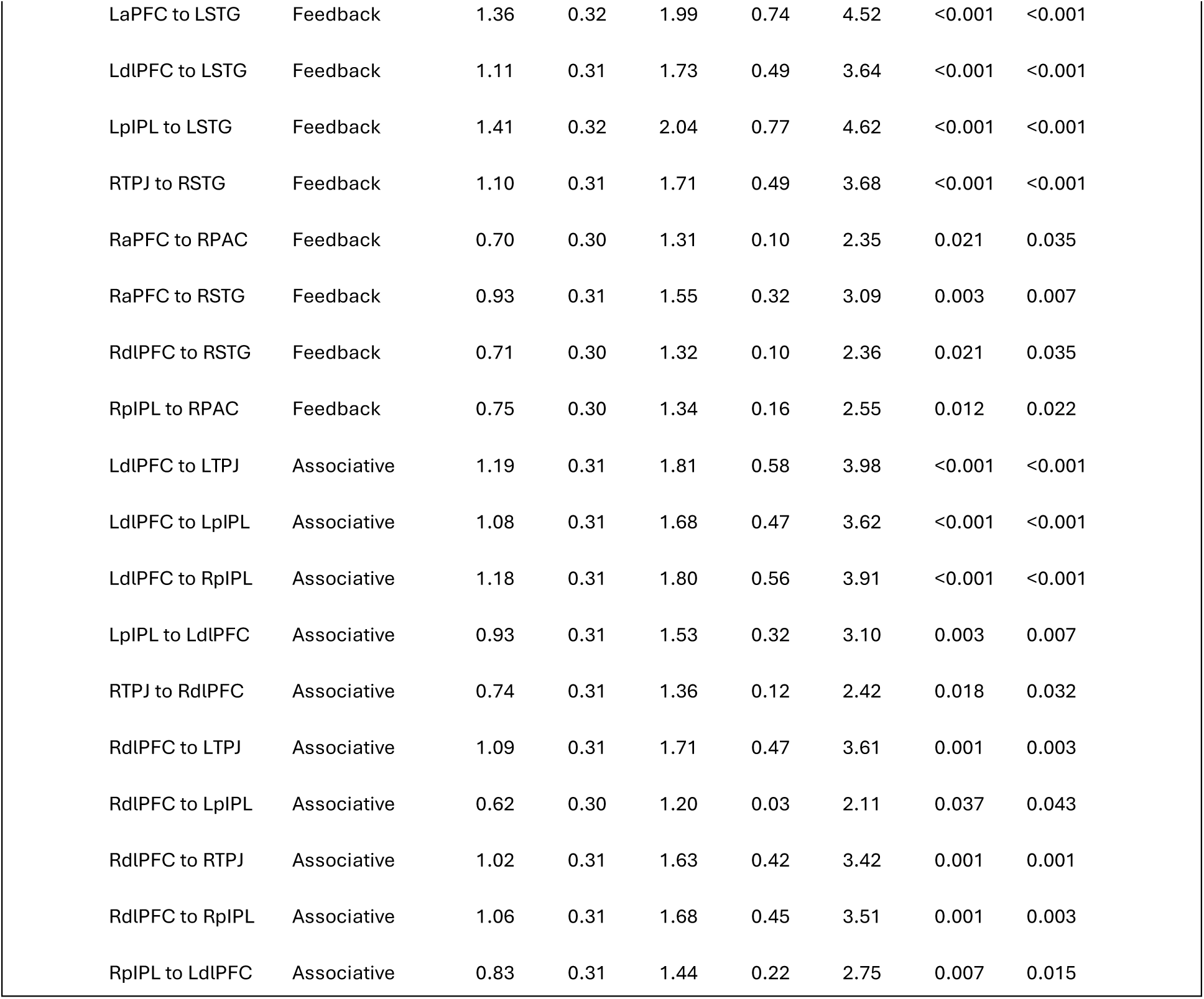
Tables 2.A.-2.D. present the results of linear mixed-effects models (LMEMs) assessing differences in directed low-gamma (37–43 Hz) connectivity between wakefulness and individual sleep stages: NREM N1, NREM N2, NREM N3, and REM. The tables report normalized estimated marginal mean differences (B), standard errors (SE), t-values, p-values (FDR-corrected), confidence intervals (95% CI), and effect sizes for each connection direction. Connections with FDR-corrected p-values < .05 are included in the tables. The normalized B estimates reflect the influence of sleep stage on connectivity strength and support direct comparisons across stages. A visual summary of these statistically significant differences is presented in Figure 1, which illustrates the spatial distribution and direction of altered connections across associative, auditory feedforward, and auditory feedback networks for each sleep stage contrast.

| Model Summary: Effects for Connections in Contrast W - N1 |  |  |  |  |  |  |  |  |
| --- | --- | --- | --- | --- | --- | --- | --- | --- |
| Effective Connectivity Analysis |  |  |  |  |  |  |  |  |
| Direction | Group | B | SE | Upper CI | Lower CI | t-ratio | p-value | p-value corr. |
| LPAC to LTPJ | Feedforward | 0.87 | 0.32 | 1.51 | 0.24 | 2.78 | 0.006 | 0.014 |
| LSTG to LTPJ | Feedforward | 1.37 | 0.35 | 2.07 | 0.68 | 4.10 | <0.001 | <0.001 |
| LSTG to LdlPFC | Feedforward | 1.25 | 0.33 | 1.91 | 0.59 | 3.91 | <0.001 | <0.001 |
| RSTG to RaPFC | Feedforward | 0.90 | 0.33 | 1.56 | 0.25 | 2.80 | 0.006 | 0.031 |
| RSTG to RdIPFC | Feedforward | 0.98 | 0.33 | 1.64 | 0.32 | 3.02 | 0.003 | 0.029 |
| LaPFC to LSTG | Feedback | 0.96 | 0.33 | 1.61 | 0.31 | 3.00 | 0.003 | 0.029 |
| LpIPL to LSTG | Feedback | 0.90 | 0.32 | 1.54 | 0.26 | 2.86 | 0.005 | 0.030 |
| RTPJ to RPAC | Feedback | 0.99 | 0.34 | 1.67 | 0.32 | 2.99 | 0.004 | 0.030 |
| RTPJ to RSTG | Feedback | 0.73 | 0.32 | 1.37 | 0.10 | 2.32 | 0.022 | 0.038 |
| LdIPFC to LTPJ | Associative | 0.98 | 0.33 | 1.63 | 0.33 | 3.05 | 0.003 | 0.014 |
| LdIPFC to LpIPL | Associative | 0.90 | 0.32 | 1.53 | 0.26 | 2.86 | 0.005 | 0.014 |
| LdIPFC to RplPL | Associative | 0.82 | 0.32 | 1.45 | 0.19 | 2.63 | 0.010 | 0.037 |
| LpIPL to LdIPFC | Associative | 0.85 | 0.32 | 1.48 | 0.23 | 2.75 | 0.007 | 0.031 |
| RdIPFC to LTPJ | Associative | 0.83 | 0.32 | 1.46 | 0.20 | 2.65 | 0.010 | 0.037 |

B.
| Model Summary: Effects for Connections in Contrast W - N2 |  |  |  |  |  |  |  |  |
| --- | --- | --- | --- | --- | --- | --- | --- | --- |
| Effective Connectivity Analysis |  |  |  |  |  |  |  |  |
| Direction | Group | B | SE | Upper CI | Lower CI | t-ratio | p-value | p-value corr. |
| LPAC to LTPJ | Feedforward | 1.58 | 0.30 | 2.18 | 0.99 | 5.66 | <0.001 | <0.001 |
| LPAC to LaPFC | Feedforward | 0.63 | 0.28 | 1.19 | 0.07 | 2.28 | 0.025 | 0.035 |
| LPAC to LdIPFC | Feedforward | 0.90 | 0.28 | 1.46 | 0.35 | 3.30 | 0.001 | 0.002 |
| LPAC to LpIPL | Feedforward | 0.96 | 0.28 | 1.52 | 0.39 | 3.45 | 0.001 | 0.002 |
| LSTG to LTPJ | Feedforward | 1.52 | 0.30 | 2.12 | 0.92 | 5.35 | <0.001 | <0.001 |
| LSTG to LdIPFC | Feedforward | 1.36 | 0.29 | 1.95 | 0.77 | 4.88 | <0.001 | <0.001 |
| LSTG to LpIPL | Feedforward | 0.91 | 0.28 | 1.47 | 0.35 | 3.32 | 0.001 | 0.002 |
| RPAC to RTPJ | Feedforward | 0.88 | 0.30 | 1.47 | 0.29 | 3.04 | 0.003 | 0.005 |
| RPAC to RdIPFC | Feedforward | 1.03 | 0.28 | 1.59 | 0.47 | 3.77 | <0.001 | <0.001 |
| RPAC to RplPL | Feedforward | 0.82 | 0.28 | 1.37 | 0.27 | 3.00 | 0.003 | 0.005 |
| RSTG to RTPJ | Feedforward | 1.15 | 0.29 | 1.72 | 0.58 | 4.17 | <0.001 | <0.001 |
| RSTG to RdIPFC | Feedforward | 0.74 | 0.28 | 1.30 | 0.19 | 2.70 | 0.008 | 0.013 |
| LTPJ to LPAC | Feedback | 0.69 | 0.28 | 1.24 | 0.14 | 2.53 | 0.013 | 0.013 |
| LTPJ to LSTG | Feedback | 0.96 | 0.28 | 1.52 | 0.40 | 3.51 | 0.001 | 0.002 |
| LaPFC to LSTG | Feedback | 1.28 | 0.29 | 1.86 | 0.70 | 4.64 | <0.001 | <0.001 |
| LdIPFC to LSTG | Feedback | 1.27 | 0.29 | 1.84 | 0.70 | 4.66 | <0.001 | <0.001 |
| LpIPL to LPAC | Feedback | 0.63 | 0.28 | 1.18 | 0.08 | 2.31 | 0.023 | 0.033 |
| LpIPL to LSTG | Feedback | 1.47 | 0.29 | 2.05 | 0.89 | 5.36 | <0.001 | <0.001 |
| RTPJ to RPAC | Feedback | 0.88 | 0.30 | 1.48 | 0.29 | 3.01 | 0.003 | 0.005 |
| RTPJ to RSTG | Feedback | 1.05 | 0.29 | 1.62 | 0.48 | 3.79 | <0.001 | <0.001 |
| RaPFC to RPAC | Feedback | 0.74 | 0.28 | 1.31 | 0.17 | 2.64 | 0.010 | 0.016 |
| RaPFC to RSTG | Feedback | 1.07 | 0.29 | 1.64 | 0.50 | 3.88 | <0.001 | <0.001 |
| RdIPFC to RSTG | Feedback | 1.35 | 0.30 | 1.94 | 0.75 | 4.75 | <0.001 | <0.001 |
| RplPL to RPAC | Feedback | 0.98 | 0.28 | 1.54 | 0.43 | 3.64 | <0.001 | <0.001 |
| LdlPFC to LTPJ | Associative | 1.16 | 0.29 | 1.73 | 0.60 | 4.24 | <0.001 | <0.001 |
| LdlPFC to LpIPL | Associative | 1.05 | 0.28 | 1.61 | 0.49 | 3.86 | <0.001 | <0.001 |
| LdlPFC to RplPL | Associative | 1.30 | 0.28 | 1.87 | 0.74 | 4.82 | <0.001 | <0.001 |
| LpIPL to LdlPFC | Associative | 1.15 | 0.28 | 1.72 | 0.59 | 4.23 | <0.001 | <0.001 |
| RdlPFC to LTPJ | Associative | 1.03 | 0.29 | 1.60 | 0.46 | 3.70 | <0.001 | <0.001 |
| RdlPFC to LpIPL | Associative | 0.72 | 0.28 | 1.27 | 0.16 | 2.61 | 0.010 | 0.012 |
| RdlPFC to RTPJ | Associative | 1.00 | 0.29 | 1.57 | 0.44 | 3.61 | <0.001 | <0.001 |
| RdlPFC to RplPL | Associative | 1.21 | 0.29 | 1.79 | 0.64 | 4.39 | <0.001 | <0.001 |
| RplPL to LdlPFC | Associative | 0.92 | 0.28 | 1.48 | 0.37 | 3.38 | 0.001 | 0.002 |
| RplPL to RdlPFC | Associative | 0.87 | 0.28 | 1.44 | 0.31 | 3.14 | 0.002 | 0.004 |

C.
| Model Summary: Effects for Connections in Contrast W – N3 |  |  |  |  |  |  |  |  |
| --- | --- | --- | --- | --- | --- | --- | --- | --- |
| Effective Connectivity Analysis |  |  |  |  |  |  |  |  |
| Direction | Group | B | SE | Upper CI | Lower CI | t-ratio | p-value | p-value corr. |
| LPAC to LTPJ | Feedforward | 1.71 | 0.30 | 2.31 | 1.10 | 6.09 | <0.001 | <0.001 |
| LPAC to LaPFC | Feedforward | 0.64 | 0.27 | 1.18 | 0.10 | 2.37 | 0.020 | 0.032 |
| LPAC to LdlPFC | Feedforward | 0.84 | 0.28 | 1.40 | 0.28 | 3.03 | 0.003 | 0.006 |
| LPAC to LpIPL | Feedforward | 0.85 | 0.28 | 1.41 | 0.29 | 3.07 | 0.003 | 0.006 |
| LSTG to LTPJ | Feedforward | 1.53 | 0.30 | 2.13 | 0.94 | 5.51 | <0.001 | <0.001 |
| LSTG to LaPFC | Feedforward | 0.62 | 0.28 | 1.18 | 0.06 | 2.24 | 0.028 | 0.041 |
| LSTG to LdlPFC | Feedforward | 1.26 | 0.29 | 1.84 | 0.68 | 4.50 | <0.001 | <0.001 |
| LSTG to LpIPL | Feedforward | 1.04 | 0.28 | 1.61 | 0.48 | 3.81 | <0.001 | <0.001 |
| RPAC to RdlPFC | Feedforward | 0.87 | 0.28 | 1.43 | 0.32 | 3.19 | 0.002 | 0.005 |
| RPAC to RplPL | Feedforward | 1.38 | 0.29 | 1.97 | 0.80 | 4.97 | <0.001 | <0.001 |
| RSTG to RTPJ | Feedforward | 0.93 | 0.28 | 1.48 | 0.37 | 3.39 | 0.001 | 0.003 |
| RSTG to RaPFC | Feedforward | 0.75 | 0.29 | 1.31 | 0.18 | 2.65 | 0.010 | 0.018 |
| RSTG to RdlPFC | Feedforward | 0.71 | 0.28 | 1.26 | 0.15 | 2.56 | 0.012 | 0.020 |
| LTPJ to LSTG | Feedback | 1.16 | 0.29 | 1.73 | 0.58 | 4.14 | <0.001 | <0.001 |
| LaPFC to LSTG | Feedback | 1.27 | 0.29 | 1.84 | 0.70 | 4.66 | <0.001 | <0.001 |
| LdlPFC to LSTG | Feedback | 1.38 | 0.29 | 1.95 | 0.80 | 5.04 | <0.001 | <0.001 |
| LpIPL to LPAC | Feedback | 0.64 | 0.28 | 1.18 | 0.09 | 2.33 | 0.022 | 0.034 |
| LpIPL to LSTG | Feedback | 1.43 | 0.29 | 2.00 | 0.85 | 5.21 | <0.001 | <0.001 |
| RTPJ to RSTG | Feedback | 1.07 | 0.29 | 1.64 | 0.50 | 3.85 | <0.001 | <0.001 |
| RaPFC to RPAC | Feedback | 0.84 | 0.28 | 1.39 | 0.28 | 3.05 | 0.003 | 0.006 |
| RaPFC to RSTG | Feedback | 1.34 | 0.29 | 1.92 | 0.76 | 4.81 | <0.001 | <0.001 |
| RdIPFC to RSTG | Feedback | 0.81 | 0.28 | 1.37 | 0.25 | 2.96 | 0.004 | 0.008 |
| RpIPL to RPAC | Feedback | 1.12 | 0.28 | 1.68 | 0.56 | 4.14 | <0.001 | <0.001 |
| LdIPFC to LTPJ | Associative | 1.32 | 0.30 | 1.91 | 0.73 | 4.66 | <0.001 | <0.001 |
| LdIPFC to LpIPL | Associative | 1.20 | 0.29 | 1.76 | 0.63 | 4.34 | <0.001 | <0.001 |
| LdIPFC to RpIPL | Associative | 0.97 | 0.28 | 1.53 | 0.41 | 3.54 | 0.001 | 0.003 |
| LpIPL to LdIPFC | Associative | 1.15 | 0.28 | 1.72 | 0.59 | 4.22 | <0.001 | <0.001 |
| RTPJ to RdIPFC | Associative | 0.62 | 0.27 | 1.17 | 0.08 | 2.29 | 0.024 | 0.036 |
| RdIPFC to LTPJ | Associative | 0.93 | 0.28 | 1.49 | 0.36 | 3.36 | 0.001 | 0.003 |
| RdIPFC to LpIPL | Associative | 0.73 | 0.27 | 1.27 | 0.18 | 2.69 | 0.008 | 0.011 |
| RdIPFC to RpIPL | Associative | 0.82 | 0.28 | 1.38 | 0.26 | 2.96 | 0.004 | 0.008 |
| RpIPL to LdIPFC | Associative | 0.70 | 0.28 | 1.25 | 0.15 | 2.56 | 0.012 | 0.020 |
| RpIPL to RdIPFC | Associative | 1.08 | 0.30 | 1.67 | 0.49 | 3.76 | <0.001 | <0.001 |

*Table 2. Tables 2.A.-2.D. present the results of linear mixed-effects models (LMEMs) assessing differences in directed low-gamma (37–43 Hz) connectivity between wakefulness and individual sleep stages: NREM N1, NREM N2, NREM N3, and REM.
| Model Summary: Effects for Connections in Contrast W – REM<br>Effective Connectivity Analysis |  |  |  |  |  |  |  |  |
| --- | --- | --- | --- | --- | --- | --- | --- | --- |
| Direction | Group | B | SE | Upper CI | Lower CI | t-ratio | p-value | p-value corr. |
| LPAC to LTPJ | Feedforward | 1.73 | 0.34 | 2.40 | 1.06 | 5.50 | <0.001 | <0.001 |
| LPAC to LdIPFC | Feedforward | 0.83 | 0.31 | 1.44 | 0.21 | 2.70 | 0.008 | 0.015 |
| LPAC to LpIPL | Feedforward | 1.05 | 0.31 | 1.67 | 0.42 | 3.43 | 0.001 | 0.003 |
| LSTG to LTPJ | Feedforward | 1.21 | 0.32 | 1.83 | 0.58 | 3.97 | <0.001 | <0.001 |
| LSTG to LaPFC | Feedforward | 1.00 | 0.32 | 1.63 | 0.37 | 3.23 | 0.002 | 0.005 |
| LSTG to LdIPFC | Feedforward | 1.00 | 0.31 | 1.62 | 0.38 | 3.29 | 0.001 | 0.003 |
| LSTG to LpIPL | Feedforward | 0.83 | 0.31 | 1.44 | 0.21 | 2.72 | 0.008 | 0.015 |
| RPAC to RdIPFC | Feedforward | 1.15 | 0.31 | 1.76 | 0.54 | 3.88 | <0.001 | <0.001 |
| RPAC to RpIPL | Feedforward | 1.03 | 0.31 | 1.64 | 0.43 | 3.47 | 0.001 | 0.003 |
| RSTG to RTPJ | Feedforward | 0.83 | 0.31 | 1.45 | 0.22 | 2.74 | 0.007 | 0.015 |
| RSTG to RdIPFC | Feedforward | 1.09 | 0.31 | 1.70 | 0.47 | 3.60 | 0.001 | 0.003 |
| LTPJ to LPAC | Feedback | 0.59 | 0.30 | 1.17 | 0.00 | 2.00 | 0.048 | 0.048 |
| LTPJ to LSTG | Feedback | 0.90 | 0.30 | 1.50 | 0.29 | 3.00 | 0.003 | 0.007 |
| LaPFC to LSTG | Feedback | 1.36 | 0.32 | 1.99 | 0.74 | 4.52 | <0.001 | <0.001 |
| LdIPFC to LSTG | Feedback | 1.11 | 0.31 | 1.73 | 0.49 | 3.64 | <0.001 | <0.001 |
| LpIPL to LSTG | Feedback | 1.41 | 0.32 | 2.04 | 0.77 | 4.62 | <0.001 | <0.001 |
| RTPJ to RSTG | Feedback | 1.10 | 0.31 | 1.71 | 0.49 | 3.68 | <0.001 | <0.001 |
| RaPFC to RPAC | Feedback | 0.70 | 0.30 | 1.31 | 0.10 | 2.35 | 0.021 | 0.035 |
| RaPFC to RSTG | Feedback | 0.93 | 0.31 | 1.55 | 0.32 | 3.09 | 0.003 | 0.007 |
| RdIPFC to RSTG | Feedback | 0.71 | 0.30 | 1.32 | 0.10 | 2.36 | 0.021 | 0.035 |
| RpIPL to RPAC | Feedback | 0.75 | 0.30 | 1.34 | 0.16 | 2.55 | 0.012 | 0.022 |
| LdIPFC to LTPJ | Associative | 1.19 | 0.31 | 1.81 | 0.58 | 3.98 | <0.001 | <0.001 |
| LdIPFC to LpIPL | Associative | 1.08 | 0.31 | 1.68 | 0.47 | 3.62 | <0.001 | <0.001 |
| LdIPFC to RpIPL | Associative | 1.18 | 0.31 | 1.80 | 0.56 | 3.91 | <0.001 | <0.001 |
| LpIPL to LdIPFC | Associative | 0.93 | 0.31 | 1.53 | 0.32 | 3.10 | 0.003 | 0.007 |
| RTPJ to RdIPFC | Associative | 0.74 | 0.31 | 1.36 | 0.12 | 2.42 | 0.018 | 0.032 |
| RdIPFC to LTPJ | Associative | 1.09 | 0.31 | 1.71 | 0.47 | 3.61 | 0.001 | 0.003 |
| RdIPFC to LpIPL | Associative | 0.62 | 0.30 | 1.20 | 0.03 | 2.11 | 0.037 | 0.043 |
| RdIPFC to RTPJ | Associative | 1.02 | 0.31 | 1.63 | 0.42 | 3.42 | 0.001 | 0.001 |
| RdIPFC to RpIPL | Associative | 1.06 | 0.31 | 1.68 | 0.45 | 3.51 | 0.001 | 0.003 |
| RpIPL to LdIPFC | Associative | 0.83 | 0.31 | 1.44 | 0.22 | 2.75 | 0.007 | 0.015 |

We compared patterns of effective connectivity in Wakefulness (W) and all the sleep stages pairwise (NREM N1, N2, N3, and REM). Overall, in all contrasts, stronger connectivity of reciprocal connections was found in Wakefulness compared to the contrasted sleep stage.

The W-N1 contrast revealed the fewest significant differences among all studied connectivity patterns. Nevertheless, the disrupted connectivity observed in the W-N1 contrast was consistently replicated in the contrasts for deeper NREM stages (W-N2 and W-N3) and with the REM stage (W-REM contrast). In the following sections we describe contrast results in more detail.

### Feedforward and Feedback connections

A detailed analysis of effective connectivity for the feedback and feedforward connections revealed a variable profile in the magnitude of changes across the W-N1, W-N2, W-N3, and W-REM contrasts. Across all contrasts, the decrease in connectivity during sleep was greater for connections of a feedforward than for feedback nature. This was reflected in higher effect sizes (as indicated by normalized *B* values), higher *t*-values, and greater statistical significance (lower *p*-values) for the feedforward connections.

For example, in the W-N1 contrast, feedforward connections, such as left STG → left TPJ and left STG → left dlPFC, showed a more robust decrease than feedback connections between these regions. Accordingly, in the W-N2 and W-N3 contrasts, feedforward pathways exhibited greater decrease of connectivity, with connections such as left PAC → left TPJ and left STG → left TPJ. Feedback connections in these contrasts remained significant but displayed lower size effects and higher variability in *p_FDR_* values. Lastly, in the W-REM contrast, feedforward connections also demonstrated a stronger reduction in connectivity (e.g., left PAC → left TPJ), than feedback connectivity.

### Associative connections

For associative connections significant decreases in connectivity during sleep were observed across all contrasts with the strongest decreases prominent in the W-N2 contrast. In the W-N1 contrast, significant decreases in directed connectivity were observed for pathways such as left dlPFC → left TPJ, left dlPFC → left pIPL, and left pIPL → left dlPFC. Importantly, across all contrasts, most associative connections showed greater decreases during NREM N2 and N3 sleep, and REM stage (W-N2, W-N3, W-REM) relative to light sleep (W-N1). In the W-N2 and W-N3 contrasts, associative connections demonstrated even larger decreases, specifically for left dlPFC → right pIPL and right dlPFC → right pIPL. In the W-REM contrast, connectivity within associative pathways were also significantly reduced compared to wakefulness, but the reduction was weaker compared to W-N2 and W-N3 contrasts, yet still stronger to decrease values in W-N1 contrast (e.g., left dlPFC → left TPJ; right dlPFC → right pIPL).

## Discussion

In this study, we investigated how fluctuations of arousal level in wakefulness and sleep modulate the directed cortical propagation of neural signals associated with 40-Hz auditory steady-state responses (ASSRs). To identify the patterns of effective connectivity underlying this propagation, we implemented the Directed Transfer Function (DTF) method in the low-gamma frequency band. Specifically, we examined which directed connections—previously implicated in auditory processing and conscious perception—showed significant reductions in strength across different sleep stages contrasted to wakefulness. To reflect their functional roles, we categorized these connections into three groups: feedforward, feedback, and associative connectivity. Our DTF analysis revealed an overall reduction in directed connectivity strength and propagation during each stage of sleep (NREM N1, N2, N3 NREM, and REM) contrasted to wakefulness. However, the pattern of these reductions was not uniform; rather, it varied in a connection-type- and sleep-stage-dependent manner, indicating a complex reorganization of cortical communication dynamics during the transition from connected to disconnected states. The strength of connections progressively weakened across NREM sleep and was particularly pronounced within auditory feedforward pathways, indicating diminished, but not abolished, efficiency of bottom-up information flow during those stages.

### Modulation of hierarchical processing of 40-Hz ASSR

We hypothesized that feedback connections between associative and auditory cortices would be most affected by changes in arousal levels. However, the results did not support this prediction. Instead, sleep-related reductions in effective connectivity were consistently more pronounced for feedforward pathways than for feedback pathways across all sleep stages.

Detailed analysis of individual connections revealed that feedforward pathways from auditory to associative regions showed more evident decreases in connectivity, supported by higher statistical effect size than feedback pathways, particularly in W-N2 and W-N3, and W-REM contrasts. While feedback connections also exhibited reductions, their degree was consistently smaller and more variable across sleep stages.

Notably, our results demonstrate a near-complete loss of connectivity between the temporal regions involved in auditory processing and associative regions over the temporoparietal junction during N2 and N3 NREM stages as compared to Wakefulness. This finding highlights the critical role of connectivity changes within the temporal and temporoparietal regions in mediating sensory processing across varying arousal states, consistent with literature emphasizing the importance of posterior cortical regions in conscious experience. The posterior parietal cortex has been implicated in integrating sensory inputs into coherent conscious awareness and supporting the subjective aspects of consciousness during dreaming and wakefulness, with disruptions in its connectivity associated with reduced consciousness during deep sleep and anesthesia (Boly et al., 2017; Koch et al., 2016; Nir & Tononi, 2010; Siclari et al., 2017; Tononi et al., 2016).

Interestingly, while auditory information partially reaches the cortex during REM sleep (Nir et al., 2015; Salinnen et al., 1996), cortical processing of auditory input remains largely limited to primary sensory regions and does not result in widespread cortical activation. These findings might support the idea of a REM sleep modulation mechanism known as ‘informational gating’; involving competition between the exogenous sensory processing and the endogenous activities, such as dream content, for computational resources (Maquet et al., 2005). This phenomenon may be reflected in the observed connectivity changes, particularly the loss of connections with higher-order regions, predominantly in the left hemisphere.

Together, these findings suggest that the reduction of 40-Hz ASSR cortical propagation during sleep is linked to changes in both feedforward and feedback connectivity, however, reduced arousal primarily disrupts bottom-up, feedforward information flow from auditory cortices to higher-order areas, rather than selectively impairing feedback signaling, as initially hypothesized.

### Reduction of associative long-range connectivity

The present findings support the hypothesis that the number of reciprocal connections between prefrontal and parietal associative regions progressively decreases with increasing NREM sleep depth. Overall, the results revealed a significant reduction in DTF values for ten connections during N3 sleep (three reciprocal), also ten connections during N2 sleep (three reciprocal), and ten connections during REM sleep (two of which were reciprocal), and only five connections during N1 sleep (one reciprocal).

Importantly, the topography of associative disconnections shifted with increasing NREM sleep depth. During N1 sleep, decreased connectivity was predominantly confined to pathways involving the left dorsolateral prefrontal cortex (dlPFC). This pattern is consistent with results previously reported by Casagrande and colleagues (Casagrande et al., 1995, 1997), who demonstrated that the left hemisphere, including frontal regions such as the dlPFC, exhibits earlier sleep onset than the right hemisphere. This hemispheric asymmetry suggests that functional disconnection of the left dlPFC may occur earlier during the transition from wakefulness to N1 sleep, potentially reflecting an adaptive mechanism of retaining some ability to detect significant changes in the environment during the sleep onset, as the right hemisphere is implied as dominant in exogenous attentional shifts (Corbetta, 1998; Spoormaker et al., 2010b; Yao et al., 2023).

During N2, N3, and REM sleep, significant decreases were also observed in pathways linking the right dlPFC to parietal associative regions. The strongest decrease in directed connectivity values across sleep stages was consistently observed for connections from the left dlPFC to both left and right pIPL. This observation is generally supported by the reports in the previous literature predicting a breakdown in cortical long-range connectivity during deep sleep (Esser et al., 2009; Lee et al., 2009; Massimini et al., 2005)

Taken together, these results indicate that reciprocal communication between prefrontal and parietal associative regions is progressively disrupted with advancing sleep depth into slow-wave NREM, as well as during REM sleep and is characterized by a reduction in bidirectional exchanges between associative regions.

In this study, we focused on the EEG activity from the same participants while they remained awake (with open eyes), and during NREM N1, N2, N3, and REM sleep stages. This has enabled us to study the ASSRs effect on physiological changes reflected by the alterations in cortico-cortical connectivity. However, here physiological activity was not compared with the reports following serial awakenings, therefore further analysis is needed that includes data on the presence or absence of sleep consciousness to investigate the effect of ASSR on the sleep experiences throughout the night.

### Conclusions

Several connectivity studies suggest that network-level mechanisms, including the posterior hot zone, frontal, and large frontoparietal networks, undergo transformations during changes of arousal. In this work, we evaluate these mechanisms using Directed Transfer Function (DTF) effective connectivity modeling on EEG data recorded during 40-Hz ASSR activity during wakefulness and sleep stages. As sleep deepens, we observed a gradual breakdown of auditory related connectivity, with NREM N1 showing relatively preserved activity in the right hemisphere, possibly reflecting sustained vigilance by attentional processes for rapid awakening to external changes. In contrast, NREM N2 and N3 sleep stages displayed the most diminished connections in auditory processing regions, particularly in the temporal and posterior parietal cortex, consistent with thalamo-cortical functional shift specific for these stages of sleep, leading to suppression of external stimulus processing at the cortical level. Contrary to our first hipothesis, feedforward connections were more disrupted than feedback ones, but in line with the second hypothesis, reciprocal prefrontal-parietal connectivity decreased with NREM depth, and showed limited recovery in REM.

The persistence of impaired connectivity from auditory to prefrontal regions in REM, despite relatively preserved associative long-range connections, supports the idea of an “informational gating” mechanism in REM sleep, which regulates the competition for processing resources between external stimuli and internally generated activities. Through functional connectivity analysis, this study advances our understanding of the neural mechanisms underlying 40-Hz auditory steady-state response processing during different sleep stages, revealing how neural processing streams are altered across states of consciousness. Furthermore, the results of this study may help to develop more effective tools for monitoring changes in patients’ consciousness, both in the context of anesthesia (e.g. assessing the degree of sedation) and severe brain injury (e.g. predicting awakening from coma, differential diagnosis of minimal consciousness and vegetative state).

## Author contributions

A.L.: conceptualization, data curation, data quality, formal analysis, visualization, writing–original draft

U.G-K.: conceptualization, data curation, data quality, investigation, project administration, resources, supervision, writing–review and editing

M.W.: data curation, formal analysis, methodology, resources, software, supervision, funding acquistion

M.H.: data curation, investigation, validation

B.Ch.: methodology, resources, and writing–original draft

M.B.: conceptualization, data curation, funding acquistion, methodology, project administration, resources, software, supervision, writing–review and editing

