## Supplemental Material for "Sleep affects low-gamma range effective cortical connectivity for 40-Hz auditory steady-state responses"

### Supplementary Material

#### Table S1. Mean DTF values across directed connectivity types

To enhance interpretability of DTF values in the results, we multiplied raw DTF values by 1000 and reported absolute mean values for each directional connection.

### Mean DTF Connectivity Values by Direction and Sleep Stage

Averaged across all participants; values reflect relative connectivity strength for each direction during distinct arousal stages.

| Direction | Arousal level | Mean DTF value |
|---|---|---|
| Associative | | |
| LdlPFC to LpIPL | N1 | 0.00148 |
| LdlPFC to LpIPL | N2 | 0.00117 |
| LdlPFC to LpIPL | N3 | 0.00088 |
| LdlPFC to LpIPL | REM | 0.00112 |
| LdlPFC to LpIPL | W | 0.00326 |
| LdlPFC to LTPJ | N1 | 0.00021 |
| LdlPFC to LTPJ | N2 | 0.00014 |
| LdlPFC to LTPJ | N3 | 0.00008 |
| LdlPFC to LTPJ | REM | 0.00013 |
| LdlPFC to LTPJ | W | 0.00059 |
| LdlPFC to RpIPL | N1 | 0.00225 |
| LdlPFC to RpIPL | N2 | 0.00113 |
| LdlPFC to RpIPL | N3 | 0.00199 |
| LdlPFC to RpIPL | REM | 0.00167 |
| LdlPFC to RpIPL | W | 0.00460 |
| LdlPFC to RTPJ | N1 | 0.00050 |
| LdlPFC to RTPJ | N2 | 0.00038 |
| LdlPFC to RTPJ | N3 | 0.00041 |
| LdlPFC to RTPJ | REM | 0.00033 |
| LdlPFC to RTPJ | W | 0.00056 |
| LpIPL to LdlPFC | N1 | 0.00268 |
| LpIPL to LdlPFC | N2 | 0.00146 |
| LpIPL to LdlPFC | N3 | 0.00150 |
| LpIPL to LdlPFC | REM | 0.00243 |
| LpIPL to LdlPFC | W | 0.00601 |
| LpIPL to RdlPFC | N1 | 0.00387 |
| LpIPL to RdlPFC | N2 | 0.00113 |
| LpIPL to RdlPFC | N3 | 0.00128 |
| LpIPL to RdlPFC | REM | 0.00086 |
| LpIPL to RdlPFC | W | 0.00221 |
| LTPJ to LdlPFC | N1 | 0.03533 |
| LTPJ to LdlPFC | N2 | 0.02516 |
| LTPJ to LdlPFC | N3 | 0.02451 |
| LTPJ to LdlPFC | REM | 0.02472 |
| LTPJ to LdlPFC | W | 0.03639 |
| LTPJ to RdlPFC | N1 | 0.01933 |
| LTPJ to RdlPFC | N2 | 0.01960 |
| LTPJ to RdlPFC | N3 | 0.02323 |
| LTPJ to RdlPFC | REM | 0.01746 |
| LTPJ to RdlPFC | W | 0.01522 |
| RdlPFC to LpIPL | N1 | 0.00309 |
| RdlPFC to LpIPL | N2 | 0.00184 |
| RdlPFC to LpIPL | N3 | 0.00181 |
| RdlPFC to LpIPL | REM | 0.00212 |
| RdlPFC to LpIPL | W | 0.00386 |
| RdlPFC to LTPJ | N1 | 0.00023 |
| RdlPFC to LTPJ | N2 | 0.00017 |
| RdlPFC to LTPJ | N3 | 0.00020 |
| RdlPFC to LTPJ | REM | 0.00015 |
| RdlPFC to LTPJ | W | 0.00048 |
| RdlPFC to RpIPL | N1 | 0.00256 |
| RdlPFC to RpIPL | N2 | 0.00142 |
| RdlPFC to RpIPL | N3 | 0.00250 |
| RdlPFC to RpIPL | REM | 0.00183 |
| RdlPFC to RpIPL | W | 0.00468 |
| RdlPFC to RTPJ | N1 | 0.00060 |
| RdlPFC to RTPJ | N2 | 0.00022 |
| RdlPFC to RTPJ | N3 | 0.00059 |
| RdlPFC to RTPJ | REM | 0.00020 |
| RdlPFC to RTPJ | W | 0.00088 |
| RpIPL to LdlPFC | N1 | 0.00334 |
| RpIPL to LdlPFC | N2 | 0.00122 |
| RpIPL to LdlPFC | N3 | 0.00186 |
| RpIPL to LdlPFC | REM | 0.00151 |
| RpIPL to LdlPFC | W | 0.00391 |
| RpIPL to RdlPFC | N1 | 0.00355 |
| RpIPL to RdlPFC | N2 | 0.00183 |
| RpIPL to RdlPFC | N3 | 0.00086 |
| RpIPL to RdlPFC | REM | 0.00240 |
| RpIPL to RdlPFC | W | 0.00407 |
| RTPJ to LdlPFC | N1 | 0.02150 |
| RTPJ to LdlPFC | N2 | 0.01364 |
| RTPJ to LdlPFC | N3 | 0.01282 |
| RTPJ to LdlPFC | REM | 0.01270 |
| RTPJ to LdlPFC | W | 0.01810 |
| RTPJ to RdlPFC | N1 | 0.01892 |
| RTPJ to RdlPFC | N2 | 0.01411 |
| RTPJ to RdlPFC | N3 | 0.00899 |
| RTPJ to RdlPFC | REM | 0.00632 |
| RTPJ to RdlPFC | W | 0.01863 |

### Mean DTF Connectivity Values by Direction and Sleep Stage

Averaged across all participants; values reflect relative connectivity strength for each direction during distinct arousal stages.

| Direction | Arousal level | Mean DTF value |
|---|---|---|
| Feedback | | |
| LaPFC to LPAC | N1 | 0.00073 |
| LaPFC to LPAC | N2 | 0.00047 |
| LaPFC to LPAC | N3 | 0.00035 |
| LaPFC to LPAC | REM | 0.00038 |
| LaPFC to LPAC | W | 0.00049 |
| LaPFC to LSTG | N1 | 0.00252 |
| LaPFC to LSTG | N2 | 0.00119 |
| LaPFC to LSTG | N3 | 0.00118 |
| LaPFC to LSTG | REM | 0.00091 |
| LaPFC to LSTG | W | 0.00627 |
| LdlPFC to LPAC | N1 | 0.00023 |
| LdlPFC to LPAC | N2 | 0.00037 |
| LdlPFC to LPAC | N3 | 0.00038 |
| LdlPFC to LPAC | REM | 0.00044 |
| LdlPFC to LPAC | W | 0.00046 |
| LdlPFC to LSTG | N1 | 0.00135 |
| LdlPFC to LSTG | N2 | 0.00055 |
| LdlPFC to LSTG | N3 | 0.00039 |
| LdlPFC to LSTG | REM | 0.00081 |
| LdlPFC to LSTG | W | 0.00248 |
| LpIPL to LPAC | N1 | 0.00042 |
| LpIPL to LPAC | N2 | 0.00028 |
| LpIPL to LPAC | N3 | 0.00028 |
| LpIPL to LPAC | REM | 0.00032 |
| LpIPL to LPAC | W | 0.00050 |
| LpIPL to LSTG | N1 | 0.00294 |
| LpIPL to LSTG | N2 | 0.00136 |
| LpIPL to LSTG | N3 | 0.00141 |
| LpIPL to LSTG | REM | 0.00171 |
| LpIPL to LSTG | W | 0.00496 |
| LTPJ to LPAC | N1 | 0.00426 |
| LTPJ to LPAC | N2 | 0.00303 |
| LTPJ to LPAC | N3 | 0.00461 |
| LTPJ to LPAC | REM | 0.00351 |
| LTPJ to LPAC | W | 0.00624 |
| LTPJ to LSTG | N1 | 0.03909 |
| LTPJ to LSTG | N2 | 0.01177 |
| LTPJ to LSTG | N3 | 0.00729 |
| LTPJ to LSTG | REM | 0.01357 |
| LTPJ to LSTG | W | 0.03207 |
| RaPFC to RPAC | N1 | 0.00039 |
| RaPFC to RPAC | N2 | 0.00024 |
| RaPFC to RPAC | N3 | 0.00021 |
| RaPFC to RPAC | REM | 0.00028 |
| RaPFC to RPAC | W | 0.00050 |
| RaPFC to RSTG | N1 | 0.00136 |
| RaPFC to RSTG | N2 | 0.00060 |
| RaPFC to RSTG | N3 | 0.00026 |
| RaPFC to RSTG | REM | 0.00077 |
| RaPFC to RSTG | W | 0.00190 |
| RdlPFC to RPAC | N1 | 0.00096 |
| RdlPFC to RPAC | N2 | 0.00117 |
| RdlPFC to RPAC | N3 | 0.00081 |
| RdlPFC to RPAC | REM | 0.00121 |
| RdlPFC to RPAC | W | 0.00126 |
| RdlPFC to RSTG | N1 | 0.00163 |
| RdlPFC to RSTG | N2 | 0.00064 |
| RdlPFC to RSTG | N3 | 0.00158 |
| RdlPFC to RSTG | REM | 0.00184 |
| RdlPFC to RSTG | W | 0.00298 |
| RpIPL to RPAC | N1 | 0.00084 |
| RpIPL to RPAC | N2 | 0.00039 |
| RpIPL to RPAC | N3 | 0.00028 |
| RpIPL to RPAC | REM | 0.00057 |
| RpIPL to RPAC | W | 0.00116 |
| RpIPL to RSTG | N1 | 0.00283 |
| RpIPL to RSTG | N2 | 0.00175 |
| RpIPL to RSTG | N3 | 0.00214 |
| RpIPL to RSTG | REM | 0.00192 |
| RpIPL to RSTG | W | 0.00296 |
| RTPJ to RPAC | N1 | 0.00163 |
| RTPJ to RPAC | N2 | 0.00187 |
| RTPJ to RPAC | N3 | 0.00357 |
| RTPJ to RPAC | REM | 0.00247 |
| RTPJ to RPAC | W | 0.00436 |
| RTPJ to RSTG | N1 | 0.01397 |
| RTPJ to RSTG | N2 | 0.00622 |
| RTPJ to RSTG | N3 | 0.00582 |
| RTPJ to RSTG | REM | 0.00503 |
| RTPJ to RSTG | W | 0.03168 |

### Mean DTF Connectivity Values by Direction and Sleep Stage

Averaged across all participants; values reflect relative connectivity strength for each direction during distinct arousal stages.

| Direction | Arousal level | Mean DTF value |
|---|---|---|
| Feedforward | | |
| LPAC to LaPFC | N1 | 0.01532 |
| LPAC to LaPFC | N2 | 0.01512 |
| LPAC to LaPFC | N3 | 0.01532 |
| LPAC to LaPFC | REM | 0.03174 |
| LPAC to LaPFC | W | 0.03312 |
| LPAC to LdlPFC | N1 | 0.07505 |
| LPAC to LdlPFC | N2 | 0.02950 |
| LPAC to LdlPFC | N3 | 0.03223 |
| LPAC to LdlPFC | REM | 0.03537 |
| LPAC to LdlPFC | W | 0.07925 |
| LPAC to LpIPL | N1 | 0.03296 |
| LPAC to LpIPL | N2 | 0.01660 |
| LPAC to LpIPL | N3 | 0.02008 |
| LPAC to LpIPL | REM | 0.01551 |
| LPAC to LpIPL | W | 0.03958 |
| LPAC to LTPJ | N1 | 0.00385 |
| LPAC to LTPJ | N2 | 0.00143 |
| LPAC to LTPJ | N3 | 0.00102 |
| LPAC to LTPJ | REM | 0.00093 |
| LPAC to LTPJ | W | 0.00682 |
| LSTG to LaPFC | N1 | 0.00359 |
| LSTG to LaPFC | N2 | 0.00237 |
| LSTG to LaPFC | N3 | 0.00235 |
| LSTG to LaPFC | REM | 0.00112 |
| LSTG to LaPFC | W | 0.00431 |
| LSTG to LdlPFC | N1 | 0.00292 |
| LSTG to LdlPFC | N2 | 0.00198 |
| LSTG to LdlPFC | N3 | 0.00274 |
| LSTG to LdlPFC | REM | 0.00444 |
| LSTG to LdlPFC | W | 0.00991 |
| LSTG to LpIPL | N1 | 0.00811 |
| LSTG to LpIPL | N2 | 0.00464 |
| LSTG to LpIPL | N3 | 0.00395 |
| LSTG to LpIPL | REM | 0.00615 |
| LSTG to LpIPL | W | 0.01254 |
| LSTG to LTPJ | N1 | 0.00036 |
| LSTG to LTPJ | N2 | 0.00023 |
| LSTG to LTPJ | N3 | 0.00022 |
| LSTG to LTPJ | REM | 0.00054 |
| LSTG to LTPJ | W | 0.00168 |
| RPAC to RaPFC | N1 | 0.05232 |
| RPAC to RaPFC | N2 | 0.02180 |
| RPAC to RaPFC | N3 | 0.01932 |
| RPAC to RaPFC | REM | 0.02802 |
| RPAC to RaPFC | W | 0.02955 |
| RPAC to RdlPFC | N1 | 0.03384 |
| RPAC to RdlPFC | N2 | 0.01948 |
| RPAC to RdlPFC | N3 | 0.02551 |
| RPAC to RdlPFC | REM | 0.01805 |
| RPAC to RdlPFC | W | 0.06200 |
| RPAC to RpIPL | N1 | 0.02139 |
| RPAC to RpIPL | N2 | 0.01935 |
| RPAC to RpIPL | N3 | 0.00725 |
| RPAC to RpIPL | REM | 0.01466 |
| RPAC to RpIPL | W | 0.03669 |
| RPAC to RTPJ | N1 | 0.00398 |
| RPAC to RTPJ | N2 | 0.00152 |
| RPAC to RTPJ | N3 | 0.00270 |
| RPAC to RTPJ | REM | 0.00247 |
| RPAC to RTPJ | W | 0.00429 |
| RSTG to RaPFC | N1 | 0.00225 |
| RSTG to RaPFC | N2 | 0.00515 |
| RSTG to RaPFC | N3 | 0.00343 |
| RSTG to RaPFC | REM | 0.00740 |
| RSTG to RaPFC | W | 0.00893 |
| RSTG to RdlPFC | N1 | 0.00270 |
| RSTG to RdlPFC | N2 | 0.00354 |
| RSTG to RdlPFC | N3 | 0.00369 |
| RSTG to RdlPFC | REM | 0.00229 |
| RSTG to RdlPFC | W | 0.00626 |
| RSTG to RpIPL | N1 | 0.00683 |
| RSTG to RpIPL | N2 | 0.00559 |
| RSTG to RpIPL | N3 | 0.00729 |
| RSTG to RpIPL | REM | 0.00889 |
| RSTG to RpIPL | W | 0.00754 |
| RSTG to RTPJ | N1 | 0.00129 |
| RSTG to RTPJ | N2 | 0.00045 |
| RSTG to RTPJ | N3 | 0.00076 |
| RSTG to RTPJ | REM | 0.00085 |
| RSTG to RTPJ | W | 0.00198 |

### Supplemental Text S1. Stimulus and preprocessing pipeline

#### B1. Sound Pressure Level calibration

To ensure the appropriate effect of the stimuli, the in-ear signal must be delivered at the correct Sound Pressure Level (SPL). Previous studies have suggested that an SPL of 65 dB is adequate for similar experiments (Neher et al., 2017), while an SPL of 60–65 dB is recommended when used in conjunction with EEG recordings (Ignatious et al., 2021). Comparable methodologies were applied in earlier work (Binder et al., 2024), where the stimuli were calibrated to 60 dB using A-weighting (dBA) to reflect levels appropriate for long-term brain stimulation studies. In the present study, A-weighted levels were also used, however, to better characterize the transient nature of the auditory stimuli, the $L_{Amax}$ metric was applied instead of $L_{Aeq}$. Given the impulsive character of the click stimuli, $L_{Amax}$ is considered more appropriate than average-based metrics, as the primary component of the signal is of impulse type (Hamernik and Hsueh, 1991; Kukulski and Wszołek, 2022).

Acoustic calibration was performed by playing stimulus using the same earphones that were used in the main study (ER-3C insert earphones, Etymotic Research, Elk Grove Village, IL, USA). Sound pressure levels were measured by in-ear microphones that were embedded in a B&K Head and Torso Simulator HATS 4128-C (Hottinger Brüel & Kjaer GmbH, Darmstadt, DE).

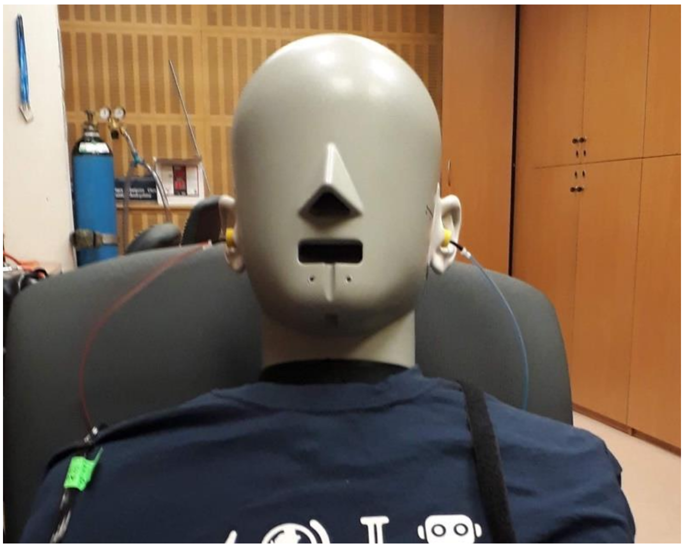

Figure S1. Calibration setup showing the B&K Head and Torso Simulator HATS 4128-C with Etymotic ER-3C insert earphones used for system calibration.

Each measurement lasted 10 seconds and was repeated iteratively until the target in-ear SPL was reached. The attenuation parameters in the measurement software were adjusted accordingly. In the repeated trials the headphones were took off the HATS ear and put again to simulate the actual behaviour while headphones are used with the human participants. After achieving the target level, at least five additional measurements were conducted to verify the stability of the stimuli. The results from these final five trials are presented in Table S3. The calibration demonstrated high stability, with a standard deviation not exceeding 0.8 dB, which is considered acceptable for ensuring the repeatability of acoustic stimulation.

Table S2. Acoustic calibration procedure results - $L_{Amax}$ from HATS after the calibration procedure for repeated measurements with the selected headphones

| Measurement no. | Left ear, dBA | Right ear, dBA |
|---|---|---|
| 1 | 59.6 | 60.4 |
| 2 | 61.0 | 60.3 |

| | | |
|---|---|---|
| 3 | 59.7 | 59.8 |
| 4 | 61.1 | 59.9 |
| 5 | 61.0 | 60.5 |
| Average | **60.5** | **60.2** |
| Std. dev. | **0.8** | **0.3** |

.

**C. Preprocessing Details**

***C.1. Sleep Scoring***

Manual sleep scoring was performed following AASM criteria, distinguishing between Wakefulness, N1, N2, N3, and REM sleep.

***C.2. Preprocessing Criteria in Atlantis Pipeline***

The Atlantis pipeline included the following steps:

- Filtering
- Artifact rejection
- Segmentation into arousal-level epochs
- Quality control at both signal and connectivity estimation levels

Only participants with data quality passing all stages were retained (see Table S1).

### Supplemental Text S2. Statistical Modeling of Directed Connectivity

To assess the impact of arousal state on directed connectivity, we used a linear mixed-effects framework across 48 cortical connections. Arousal was categorized as: Wakefulness (reference), N1, N2, N3, and REM.

Model specification:

lmer(value ~ arousal_level + (1 | id), data = directed_connectivity, control = lmerControl(optimizer = "bobyqa"))

Post-estimation:

- Estimated marginal means: emmeans package
- Effect sizes: Cohen's d via effectsize package
- P-values adjusted for multiple comparisons using the Benjamini–Hochberg FDR method

Singularity Warnings: 8/48 models produced warnings, indicating low variance or overfitting.

Reference: https://decision-lab.org/wp-content/uploads/2020/07/SOP_Mixed_Models_D2P2_v1_0_0.pdf

Despite convergence, 8 directional models yielded singularity warnings. We verified no data import errors or arousal-level imbalances. After applying the "bobyqa" optimizer, only RpIPL → RPAC resolved. For remaining models, we removed the random intercept and compared full vs. simplified models via AIC/BIC. Simplified fixed-effects models were retained when they performed equally or better.

Problematic connections:

- Feedforward: LPAC → LTPJ
- Feedback: RTPJ → RSTG, LTPJ → LPAC, RpIPL → RPAC
- Associative: LdlPFC → LTPJ, LdlPFC → LpIPL, RdlPFC → RTPJ, RdlPFC → LpIPL

Model Adjustment:

Simplified versions (fixed-effect models using lm()) were estimated and compared via AIC and BIC. Fixed-effect models provided comparable or better fit and were retained for final interpretation.

### Supplemental Text S3. Connectivity Type x Arousal Interaction

To investigate how connectivity patterns differed by sleep stage and direction type, we performed two-way ANOVAs using:

value ~ sleep * connectivity_type

Contrast coding:
- Associative: 1/0/0
- Feedforward: 0/1/0
- Feedback: 0/0/1

Post hoc analysis:
- Pairwise comparisons using emmeans
- FDR-adjusted p-values
- Cohen's d calculated for feedforward and feedback connections between each sleep stage and wakefulness

Reporting:
All results reported using gt package (R) in publication-ready tables. Significance threshold: $p < .05$
